# COVID-19: Salient Aspects of Coronavirus Infection, Vaccines and Vaccination Testing and their Implications

**DOI:** 10.1101/2021.12.21.470882

**Authors:** Pradeep K. Pasricha, Arun K. Upadhayaya

## Abstract

In the present study, three basic aspects related to COVID-19 are presented.

a. The occurrence of coronavirus infection is analyzed statistically as number of coronaviruses infected alveolar cells compared to normal alveolar cells in human lungs. The mole concept is used to estimate the number of normal alveolar cells per human lung. The number of coronavirus infections in infected alveolar cells is estimated from the published Lower Respiratory Tract (LRT) load data. The Poisson probability distribution is aptly applied to imply the incubation period of the coronavirus infection to be within day-3 to day-7, with the cumulative probability of 75%. The incubation period within day-0 to day-10 has a cumulative probability of 98%. It implies a 10-day quarantine to isolate an uninfected individual as a precautionary measure.
b. Three vaccines to combat COVID-19, which adopt distinct paradigms while preparing them, are analyzed. These are Moderna’s mRNA-1273, Oxford-AstraZeneca’s ChAdOx1 nCoV-19 and Bharat BioTech’s COVAXIN. The mole concept is used to estimate the antigen mass density per dose of each of these vaccines as 10 g cm^-3^, 0.1 g cm^-3^ and 1 g cm^-3^, respectively. The vaccines are deemed to be compatible to neutralize the infection.
c. A statistical analysis is performed of the Moderna’s mRNA-1273 vaccine efficacy of 94.1% and Oxford’s ChAdOx1 nCoV-19 vaccine efficacy of 62.1% in terms of groups of volunteers testing negative to vaccine by chance. In the Moderna vaccination testing scenario, since the probability of negative response of vaccine is small, the Poisson probability distribution for 95% cumulative probability is used to describe the vaccination testing in 300 samples of 47 volunteers each. Thus, 87% of samples have average group of 3 volunteers testing negative to vaccine. About 6% of samples have all volunteers testing positive to vaccine. In the Oxford vaccination testing scenario, since the probability of negative response of vaccine is finite, the Gaussian probability distribution for 95% probability is used to describe the vaccination testing in 75 samples of 120 volunteers each. Thus, 68% of samples have average group of 45 volunteers testing negative to vaccine. No sample has all volunteers testing positive to vaccine. A vaccine, irrespective of its efficacy being high or low, is necessary for mass immunization.

## 1. Introduction

In the present study, three basic aspects related to COVID-19 are presented. These are (i) the statistical analysis of the occurrence of coronavirus infection in human beings, (ii) the analysis of varied vaccines in terms of antigen mass density per dose to neutralize the infection and (iii) the statistical analysis of a vaccine testing procedure in terms of vaccine efficacy. In (i) and (ii), the mole concept, that is, one mole of a substance (chemical, biological, virus, or vaccine) contains Avogadro’s number of particles (atoms, molecules, cells, viral particles, or specific antigens) is applied.

i. A statistical study is made for the appropriate probability distribution to describe the occurrence of coronavirus infection in terms of the number of coronavirus infected alveolar cells vis-à-vis normal alveolar cells in human lung. The number of normal alveolar cells in the human lung is evaluated through the mole concept. The number of infected alveolar cells is estimated from the published Lower Respiratory Tract (LRT, moderate to severe COVID-19 symptoms) viral load data in fourteen patients from Germany and one patient from China. The number of coronavirus infections in infected alveolar cells (per lung volume) is much less than the number of normal alveolar cells (per lung volume). Thus, the Poisson probability distribution may be applied to analyze the probability of occurrence of coronavirus infection in human beings.
ii. Three vaccines used to combat COVID-19, which adopt distinct paradigms while preparing them, are analyzed. These are Bharat BioTech’s COVAXIN, Oxford-AstraZeneca’s ChAdOx1 nCoV-19 (AZD1222) and Moderna’s mRNA-1273. The vaccine COVAXIN is prepared using live but inactivated SARS-CoV-2 virus. The vaccine AZD1222 is a genetically modified preparation from chimpanzee adeno virus. The vaccine mRNA-1273 is a messenger (m) RNA-based vaccine that carries the SARS CoV-2 virus spike immunogen. The mole concept is used to estimate the antigen mass density per dose of each of these vaccines in order to make a comparative study amongst these vaccines.
iii. A pertinent parameter to gauge the occurrence and severity of COVID-19 is immunity amongst masses. Consequently, in the so-called Phase 3 of vaccination testing, a large number of volunteers, as finite numbers of volunteers at a number of centers, are tested for vaccine efficacy. In mathematical statistics, a large number of volunteers form a population. The finite numbers of volunteers (at each center) are the samples of the population. A probability distribution of the negative response of volunteers to vaccine in samples is thus evaluated; which enables to determine the vaccine efficacy. A number of vaccines for the COVID-19 pandemic have been developed, with a range of mean vaccine efficacy of 60% to 95%. The outcome of vaccination testing of Moderna’s mRNA-1273 vaccine and Oxford’s ChAdOx1 nCoV-19 vaccine in terms of the respective vaccine efficacy are analyzed.

The Moderna’s mRNA-1273 vaccine, co-developed by Moderna, Inc., and NIAID (National Institute of Allergy and Infectious Diseases), was administered to 14134 volunteers at 99 centers in Phase 3 in the USA [2, 3, 4]. (The volunteers received two doses of the vaccine.) The volunteers were further grouped into three categories, (a) persons 65 years of age or older, (b) persons younger than 65 years of age who are at risk for severe COVID-19 and (c) persons younger than 65 years of age not at risk for COVID-19. There were subgroups based on gender, etcetera. In the statistical analysis of the sample (comprising subsamples), the data on the number of volunteers affected by COVID-19 are treated as random. The mean vaccine efficacy is 94.1% at preventing symptomatic COVID-19. The 95% confidence interval (or confidence limits) for the vaccine efficacy is 89.3% - 96.8% of the Poisson probability distribution of the sample data (number of affected volunteers, efficacy). These are the range of values of the population mean efficacy, calculated using the sample mean efficacy, with 95% certainty. In the present study, since the probability of negative response of vaccine is small, the Poisson probability distribution is used to describe the vaccine testing procedure.

The Oxford’s ChAdOx1 nCoV-19 vaccine, developed at Oxford University, was administered to 8895 volunteers at 25 centers in Phase 2 and Phase 3 in the UK and Brazil. (The vaccine is popularly known as Oxford-AstraZeneca COVD-19 vaccine AZD1222. The volunteers received two standard doses (SD, SD/SD) of the vaccine. The vaccine was also administered to volunteers in South Africa at a number of centers. However, the results of the statistical analysis of the vaccine trials in South Africa have not been reported.) The volunteers were further grouped into three categories, (a) persons 70 years of age or older, (b) persons in the age group 56 - 69 years and (c) persons in the age group 18 - 55 years. There were subgroups based on gender, etcetera. The statistical analysis of the efficacy in the randomized subsamples of volunteers at risk for COVID-19 was affected though a robust Poisson regression model. The mean vaccine efficacy is 62.1% at preventing symptomatic COVID-19. The 95% confidence interval (or confidence limits) for the vaccine efficacy is 41.0% - 75.7% for the determination of the mean efficacy with 95% certainty. In the present study, since the probability of negative response of vaccine is finite, the Gaussian probability distribution is used to describe the vaccine testing procedure.

## 2. Probability Distribution of Occurrence of Coronavirus Infection

The Poisson probability distribution is aptly applied to infer the incubation period of the coronavirus infection in human beings.

### 2.1 Number of Alveolar Cells in Human Lung

One may estimate the number of alveolar cells in the human lung as follows. The molecular mass (weight) of biological molecules, such as proteins, in lung tissue cells, is taken as M_w_ ≈ 10^5^ Da (dalton, g mol^-1^, grams mole^-1^) [1, Table 12.1]. The mass density of lung tissue is ρ ≈ 0.3 g cm^-3^ [2]. The average adult lung volume is 1.5 litres (1.5 × 10^3^ cm^3^). (The lung volume is the volume of air remaining in lungs after a normal exhalation, called the functional residual capacity.)

Molecular mass M_w_ = 10^5^ Da implies that 1 mole of lung tissue has a mass of 10^5^ g.1 cm^3^ of lung tissue has ρ/M_w_ = 0.3/10^5^ = 3 × 10^−6^ moles.

1 mole contains 6.02 × 10^23^ cells of lung tissue. (The Avogadro constant, N_A_ = 6.02 × 10^23^ mol^-1^, is the number of atoms, molecules, or cells.)

So, N = N_A_ x moles = N_A_ × ρ/M_w_ = 6.02 × 10^23^ × 3 × 10^−6^ = 1.806 × 10^18^ cells occupy a volume of 1 cm^3^.

Volume V (cm^3^) per cell = 1 / N = (1 / N_A_) x M_w_ / ρ = 1 / (1.806 × 10^18^) = 5.537 × 10^−19^ cm^3^.

The volume occupied per cell is of the order of d^3^, where d is the size parameter of a cell. So, approximately, d = V^1/3^ = (5.537 × 10^−19^)^1/3^ ≈ 0.821 × 10^−6^ cm ≈ 8.21 nm (nanometer, 1 nm = 10^−9^ m). Hence, 1.806 × 10^18^ numbers of cells of size 8.21 nm occupy a volume of 1 cm^3^.

The “ready to use” expression of volume V (nm^3^) of a substance (chemical, biological, virus, or vaccine) in terms of molecular mass M_w_ (Da) and density ρ (g cm^-3^) is given as [1]

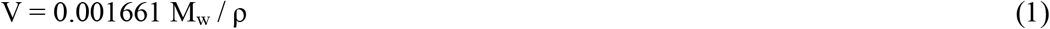

where 0.001661 is (1/6.02) × 10^−2^. 6.02 is part of the Avogadro number.

Thus, the size parameter d (nm) is

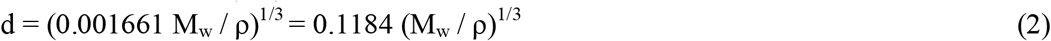

Also, the density ρ (g cm^-3^) is

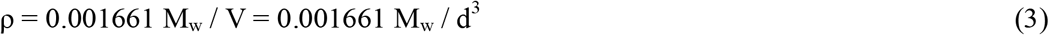

The alveoli, minute air sacs in the lungs, provide an enormous surface area for efficient respiration. The tissue cells of alveoli, the alveolar macrophages, are large scavenger cells. They remove foreign bodies such as dust, viruses and bacteria from blood, tissues and organs. In essence, the immune cells, B cells and T cells, types of “small” white blood cells, act on viruses. The T cells “memorize”, “identify” and “remember” how to fight the virus infection. The B cells “make” antibodies to neutralize the virus infection before the viruses infect cells. The virus infected “perished” cells are also removed in the process. The size of B cells and T cells, which are in globular shape, is: d ≈ 8000 nm [1, Table 12.2]. The alveolar cell size is taken as d ≈ 8000 nm. One may estimate the number of alveolar cells of size parameter 8000 nm in terms of cells of 8.21 nm in a volume of 1 cm^3^ as: 1.806 × 10^18^ × (8.21)^3^ / (8000)^3^ ≈ 2 × 10^9^ cells. The “scaling factor” to transform cells of size parameter 8.21 nm to cells of 8000 nm is: (8.21)^3^ / (8000)^3^ ≈ 1.08 × 10^−9^ ≈ 10^−9^. Since the average lung volume is about 1.5 × 10^3^ cm^3^, the estimated number of cells of size parameter 8.21 nm is: 1.5 × 10^3^ × 1.806 × 10^18^ ≈ 2.7 ×10^21^ ≈ 10^21^. Also, the number of alveolar cells of size parameter 8000 nm is: 1.5 × 10^3^ × 2 ×10^9^ ≈ 3.0 × 10^12^ ≈ 10^12^. The number of alveolar cells is given as 10^10^ × 10^±1^ [3].

### 2.2 Number of Coronavirus Infected Alveolar Cells in Human Lung

The alveolar cells are the host cells for the replication of coronavirus in the human lung. Normally a human does the breathing process of inhalation and exhalation of air about 16 times a minute, about half a litre (500 ml or cm^3^) of air each time called the tidal volume. Not all of that half liter breath reaches the alveoli, the minute air sacs in the lungs. One-third of it shuffles in and out of the windpipe and other passages.

A coronavirus penetrates a cell. It replicates to produce virus particles, the virions. Overwhelmed by the virions, the cell perishes. Then the released virus and virions, collectively the viruses, attack other cells. It is anticipated that one is infected by coronavirus in about ten minutes after close contact with an infected person. It is termed the virion entry into cell [3]. Thus, the virus-infected volume of air that could infect the human lung in ten minutes is about: 10 × 1/3 × 16 × 500 cm^3^ ≈ 2.7 ×10^4^ cm^3^ ≈ 10^4^ cm^3^, which is greater than the lung volume of 1.5 × 10^3^ cm^3^. However, the initial virus dose does not decide the severity of infection, that is, the number of infected alveolar cells. It is the activation of an individual’s immune system, triggered by the coronavirus, which decides the number of infected alveolar cells. Also, the immune response (immunity) of the human body, in terms of white blood cells (B cells and T-cells) and the presence of antibodies against the virus, varies with individuals. A measure of “healthy” immunity in terms of the amount (mass density, number density) of immune B- and T- cells is unknown. The amount of B- and T-cells is a function of many factors, including age [4]. The number of infected alveolar cells may be estimated in terms of the viral load of an infected individual. The viral load is measured as the quantity of virus “copies” per unit volume (millilitre^-1^, ml^-1^, or cm^-3^). The viral load data has been reported in fourteen patients from Germany and one patient from China [5]. Two sets of data are given, one each for Upper Respiratory Tract (URT, mild COVID-19 symptoms) and Lower Respiratory Tract (LRT, moderate to severe COVID-19 symptoms, lung infection). (The mild URT symptoms may include fever, without shortness of breath, that is, normal respiratory rate. The moderate LRT symptoms may include breathlessness, high respiratory rate and low blood oxygen levels. The severe LRT symptoms may include breathlessness, higher respiratory rate and lower blood oxygen levels.) In general, the URT virus load is lower than the LRT viral load by two orders of magnitude, that is, by a factor of 100 (10^2^) [5]. The LRT virus load may be used to estimate the infected alveolar cells as follows. The LRT viral load of a “representative” patient#3 in Germany is about 10^6^ copies ml^-1^ on day-4 and day-5 of infection. The viral load is about 10^5^ copies ml^-1^ from day-6 to day-15 of infection. The viral load “exponentially” falls to about 10^4^ copies ml^-1^ from day-16 to day-20. Here, the minimum detectable viral load is 100 (10^2^) copies ml^-1^. The LRT viral load of the patient in China is about 10^5^ copies ml^-1^ and 10^7^ copies ml^-1^ on day-3 and day-4– to – day-6 of infection, respectively. Then the viral load “exponentially” falls to about 10^5^ copies ml^-1^ on day-8 and about 10^3^ copies ml^-1^ on day-9 – to – day-11 of infection, respectively. Here, the minimum detectable viral load is 1 (10^0^) copies ml^-1^. It seems reasonable to assume a viral load of 10^6^ copies ml^-1^ (on day-5) to estimate the number of infected alveolar cells. The average number of virions produced by a single infected cell, the “per cell viral yield”, termed Burst size, is 10^3^ virions [3]. The coronavirus size parameter is 100 nm [3]. Thus, the number of coronavirus infected alveolar cells is: 10^6^/10^3^ ml^-1^ = 10^3^ ml^-1^ (or cm^-3^). Since the average lung volume is about 1.5 × 10^3^ cm^3^, the estimated number of infected alveolar cells in the human lung is: 1.5 ×10^3^ × 10^3^ ≈ 1.5 × 10^6^ ≈ 10^6^.

### 2.3 Poisson Probability Distribution of Coronavirus Infection

The probability of occurrence of coronavirus infection is small. The number of infections in terms of the infected alveolar cells of 10^3^ (per lung volume, Section 3) is much less than the number of alveolar cells of 10^9^ (per lung volume, Section 2). A Poisson distribution function is an analytical form for the probability distribution of such data [6, Chapter 3]. The Poisson probability distribution is applicable when the probability of an event is very small, small, or unknown. For illustration, from Section 3, the LRT viral load per day, over a period of 15 days, are the set of infection events. In the Poisson distribution function, day-5 is roughly the mean of days of infection events, corresponding to the maximum of LRT viral load (on day-5). (The model fitted curves in Fig. 2 and Fig. 4B in [5] are bounded essentially only on the left side of the mean, which is considered appropriate for the Poisson probability distribution [6, Chapter 3]). With mean µ = 5 day, one may compute the probability of an infection on day-0, day-1, day-2,…, day-15 of the occurrence of coronavirus infection. Hence, these probabilities may be multiplied by the viral load of 10^6^ copies ml^-1^ to estimate the viral load and the infected alveolar cells on each day. The maximum LRT viral load is on day-5 of the 15-day infection period, for example. For the Poisson distribution of LRT viral load with day, given by the variable X, the mean µ is considered significant within ±σ. One has X = µ ± σ, where σ is the standard deviation. Here, µ = 5 and 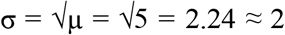 (day). Thus, X = 5 ± 2; or, 3 ≤ X ≤ 7. The infection symptoms may appear within day-3 to day-7 of the exposure to infection, mean day-5 (µ = 5), with a 75% cumulative probability of infections. The range of occurrence of infections 3 ≤ X ≤ 7 is the incubation period [5]. For the mean µ to be significant within ±2σ, X = µ ± 2σ = 5 + 2 × 2.24 ≈ 5 ± 5 (0 ≤ X ≤ 10), the cumulative probability for a Poisson distribution with mean day-5 (µ = 5) is 98%. Thus, the cumulative probability of the occurrence of infections is 98% for the exposure to infection within day-0 to day-10.

## 3. COVID-19 Vaccines

A COVID-19 vaccine uses a specific antigen, which stimulates in the human body the development of antibodies through B cells and the response of T cells to neutralize the effect of the antigen. It thereby confers immunity against the COVID-19. Three vaccines are analyzed that adopt distinct paradigms while preparing them. In particular, the antigen mass density per dose of each of these vaccines is calculated. Estimation is made of the mass density of SARS-CoV-2 antibody secreted by B cells. Also, the mass density of B cells, T cells and antibodies in the human beings is estimated. In an inert solution, the injection dose in the intramuscular/intravenous vaccination is 0.5 ml in each dose for these vaccines.

The Bharat BioTech vaccine, COVAXIN, is prepared using live but inactivated SARS-CoV-2 virus (antigen per dose of mass 6 µg, intramuscular injection dose in a potassium buffer solution of 0.5 ml and vaccine efficacy 81 %. COVAXIN is co-developed by the biotechnology company Bharat Biotech and the National Institute of Virology (NIV) of the Indian Council of Medical Research (ICMR) [7].) The molecular mass of SARS-CoV-2 virus, size (d) ∼ 100 nm, density (ρ) ∼ 1 g cm^-1^, is M_w_ ∼ 10^9^ Da [3]. Thence, the molecular mass of SARS-CoV-2 antigen is taken as ∼ 10^9^ Da.

From (3), the density (ρ) of virus = 0.001661 × 10^9^ / 100^3^ ≈ 1.66 g cm^-3^ ∼ 1 g cm^-3^.

1 mole of antigen has a mass of 10^9^ g, and has 6.02 × 10^23^ inactivated viruses.

So, 6 µg of antigen has: 6 × 10^−6^ / 10^9^ = 6 × 10^−15^ mole.

Also, 6 µg has: 6.02 × 10^23^ × 6 × 10^−15^ = 3.6 × 10^9^ ≈ 3 × 10^9^ inactivated viruses.

Thus a COVAXIN dose, mass 6 µg, molecular mass 10^9^ Da, has 3 × 10^9^ antigen particles.

The Oxford-AstraZeneca vaccine, ChAdOx1 nCoV-19, AZD1222, is a genetically modified preparation from chimpanzee adenovirus (viral particles per dose of 5 × 10^10^, intravenous injection dose in an inert solution of 0.5 ml and vaccine efficacy 62.1 %. AZD1222 is developed at Oxford University [8].) The molecular mass of Chimpanzee adenovirus, size (d) ∼ 100 nm, is M_w_ ∼ 150 × 10^6^ Da (= 1.5 x10^8^ Da) [9].

A dose of AZD1222 has mass = 1.5 ×10^8^ × 5 × 10^10^/ 6.02 × 10^23^ ≈ 1.2 × 10^−5^ g ≈ 10 µg.

Thus an AZD1222 dose, mass 10 µg, molecular mass 1.5 × 10^8^ Da, has 5 ×10^10^ antigen particles.

The Moderna vaccine, mRNA-1273, is a messenger (m) RNA-based vaccine that carries the SARS CoV-2 virus spike immunogen (antigen per dose of 100 µg, intramuscular injection dose in an inert solution of 0.5 ml and vaccine efficacy 94.1 %. mRNA-1273 is co-developed by Moderna, Inc., and NIAID (National Institute of Allergy and Infectious Diseases) [10, 11, 12].) The molecular mass of mRNA molecule, a nanowire of radius (r) ∼ 1 nm and linear length (h) ∼ 300 nm, volume (V, π r^2^h) ≈ 900 nm^3^, is M_w_ ∼ 6 × 10^6^ Da [13].

So, 100 µg of antigen has: 6.02 × 10^23^ × 100 × 10^−6^/ 6 ×10^6^ = 10^13^ antigen particles.

Thus an mRNA-1273, mass 100 µg, molecular mass 6 × 10^6^ Da, has 10^13^ antigen particles.

From (3), the density of a COVAXIN antigen dose is: 0.001661 × 10^9^ / 100^3^ ≈ 1.6 g cm^-3^. Also, the density of an AZD1222 antigen dose is: 0.001661 × 1.5 × 10^8^ / 100^3^ ≈ 0.3 g cm^-3^. Also, the density of an mRNA-1273 antigen dose is: 0.001661 × 6 × 10^6^ / 900 ≈ 10 g cm^-3^. These are the estimated values of density of an antigen dose of vaccines. It is concluded that the density of an antigen dose is ∼ 10^0±1^ g cm^-3^. It is instructive to note that the density of normal human red blood (cells) and that of white blood cells are ∼ 1.1 g cm^-3^ (ml). (Normally, the white blood cells density (≈ 1.08 g cm^-3^) is lower than the red blood cell density (≈ 1.1 g cm^-3^.)

The mass density of SARS-CoV-2 antibody, secreted by B cells, may be estimated as follows. The molecular mass of SARS-CoV-2 antibody is ∼ 114 × 10^3^ Da [14]. The “effective” size of antibody, which is “heavier” Y-shaped molecule, is: d ≈ 10 nm [15]. Thus, from (3), the mass density of SARS-CoV-2 antibody is: 0.001661 × 114 × 10^3^ / 10^3^ ≈ 0.2 g cm^-3^. An estimate of molecular mass of antibodies, secreted by B cells, is: 150 × 10^3^ Da [15]. Thus, the mass density of antibodies is: 0.001661 × 150 × 10^3^ / 10^3^ ≈ 0.25 g cm^-3^. The mass density of SARS-CoV-2 antibody is comparable to the mass density of antibodies secreted by B cells. It is worthwhile to estimate the mass density of B and T cells in white blood cells. The molecular mass of B and T cells is ∼ 70 × 10^3^ Da [16]. From (3), with d ≈ 8000 nm, the mass density of B and T cells is: 0.001661 × 70 × 10^3^ / 8000^3^ ≈ 0.2 × 10^−9^ g cm^-3^ (0.2 ng cm^-3^, nanogram cm^-3^).

## 4. COVID-19 Vaccination Testing

The Poisson probability distribution is applied to analyze Moderna’s mRNA-1273 vaccine for 94.1% vaccine efficacy. The Gaussian probability distribution is applied to analyze Oxford’s ChAdOx1 nCoV-19 vaccine for 62.1% vaccine efficacy. Thereby, the percentage of samples of groups of volunteers testing negative to vaccine is determined for 95% cumulative probability and 95% probability of the Moderna’s vaccine and Oxford’s vaccine, respectively.

### 4.1 Statistical Analysis of Moderna’s 94.1% Vaccine Efficacy

In Phase 3, in Moderna’s vaccine testing procedure, there are 14134 volunteers at effectively 297 (3 × 99 = 297) centers at an average of 47.59 volunteers per center [10, 11, 12]. (The number of groups of persons based on the age criterion is 3 at 99 centers.) The present study considers the vaccine testing in a population of 14100 volunteers, as N = 300 samples of n = 47 volunteers each. The vaccine efficacy is 94.1%. Thus, the probability (p) of negative response of vaccine in a sample is 5.9%; p = 0.059. Also, the average number of volunteers (µ) in a sample testing negative to vaccine is: µ = n × p = 47 × 0.059 = 2.773 (≈ 3). Since p << 1 and, therefore, µ << n, the Poisson probability distribution aptly describes the vaccine testing procedure. The Poisson distribution function, with mean equal to 2.773 (µ) volunteers testing negative to the vaccine, is used to compute the probability of groups of volunteers (say, a group of 3 volunteers) testing negative to the vaccine in a sample. Hence, by chance, any groups of volunteers in 300 samples testing negative to vaccine are determined. These groups of volunteers testing negative for 95% confidence interval are examined vis-à-vis the average number of volunteers obtained with 89.3% and 96.8% vaccine efficacy (95% confidence interval 89.3% - 96.8%).

The Poisson probability distribution is appropriate for describing small samples of a large population [6, Chapter 3]. Here, a sample means a finite number of volunteers being tested for vaccine, that is, 47 volunteers. A population refers to a large number of volunteers being tested for vaccine, that is, 14100 volunteers – 300 samples of 47 volunteers each.

The Poisson distribution function is given by

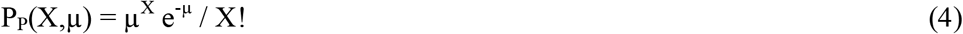

where P_P_(X,µ) is the “discrete” probability of X number of volunteers in a sample (per sample) testing negative to vaccine (given by the discrete random variable X), µ the mean (average) number of volunteers per sample testing negative to vaccine and ∑_x_P_P_(X,µ) = 1. Here, the probability (p) of a volunteer testing negative to vaccine is 100% - 94.1% = 5.9%, that is, p = 0.059. A sample has n = 47 volunteers. So, µ = np = 47 × 0.059 = 2.773, which is the average number (≈ 3) of volunteers testing negative to vaccine in a sample (per sample). Given, total number of samples of volunteers N = 300. The sign “!” means the factorial, X! = X x (X-1) x (X-2) x (X-3)…; 0! = 1. N x P_P_(X,µ) is the number of samples having X number of volunteers testing negative to vaccine, by chance.

The number of samples (m) having no volunteer testing negative is given by X = 0,

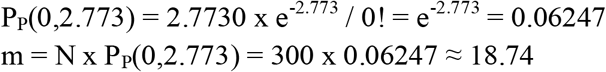

Thus, 18.74 samples have no volunteer testing negative, all 47 volunteers testing positive, by chance. The number of volunteers who test positive is: 18.74 × 47 ≈ 881.

In an analogous manner, one may compute the number of samples (m) of volunteers testing negative for each of variable X = 1, …, X = 12, with a constant C = P_P_(0,2.773) = 0.06247.

For example, the number of samples (m) having one volunteer testing negative is given by X = 1,

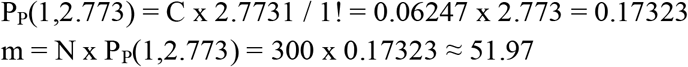

Thus, 51.97 samples have 1 volunteer testing negative, 46 volunteers testing positive, by chance. The number of volunteers who test negative is 52 (≈ 51.97 × 1). The number of volunteers who test positive is 2391 (≈ 51.97 × 46). Total number of volunteers is 2443 (= 52 + 2391).

The number of samples (m) having two volunteers testing negative is given by X = 2,

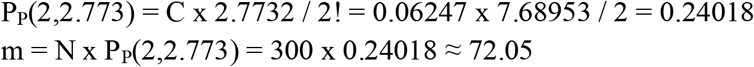

Thus, 72.05 samples have 2 volunteers testing negative, 45 volunteers testing positive, by chance. The number of volunteers who test negative is 144 (≈ 72.05 × 2). The number of volunteers who test positive is 3242 (≈ 72.05 × 47). Total number of volunteers is 3386 (= 144 + 3242).

A summary of number of samples of volunteers and number of volunteers therein testing negative (and positive) to vaccine, computed using expression (4), is given in Table 1. The variable X is a measure of number of volunteers in a sample (per sample) testing negative to vaccine. P_P_(X,µ) is the corresponding probability. The tabulated numbers of samples of volunteers testing negative (m) are bounded on the left side of the mean of 2-3 (µ) volunteers testing negative, by chance, and exhibit Poisson probability distribution of variable X. The number of samples of volunteers testing negative is 299.99 (≈ 300, N). The average number of volunteers testing negative is 832, p x 14100 = 0.059 × 14100 = 831.9 ≈ 832, in the population of 14100. Alternately, one has: µ x 300 = 2.773 × 300 = 831.9 ≈ 832 volunteers testing negative. The average number of volunteers testing negative per sample of 47 (n) volunteers is 2.773 (≈ 3, µ).

**Table 1.** Summary of number of samples and volunteers, in 300 samples of 47 volunteers each, testing negative (and positive) for 94.1% vaccine efficacy, by chance (average number of volunteers testing negative = 832)

| | Variable (X) | Probability<br>$P_P(X, 2.773)$ | Cumulative<br>probability<br>$\sum P_P(X, 2.773)$ | Number of<br>samples<br>(m) | Number of<br>volunteers<br>testing<br>negative | Number of<br>volunteers<br>testing<br>positive | Number of<br>volunteers |
| --- | --- | --- | --- | --- | --- | --- | --- |
|  | 0 | 0.06247 | 0.06247 | 18.74 | 0 | 881 | 881 |
|  | 1 | 0.17323 | 0.23570 | 51.97 | 52 | 2391 | 2443 |
|  | 2 | 0.24018 | 0.47588 | 72.05 | 144 | 3242 | 3386 |
|  | 3 | 0.22201 | 0.69789 | 66.60 | 200 | 2930 | 3130 |
|  | 4 | 0.15391 | 0.85180 | 46.17 | 185 | 1985 | 2170 |
|  | 5 | 0.08536 | 0.93716 | 25.61 | 128 | 1076 | 1204 |
|  | 6 | 0.03945 | 0.97661 | 11.84 | 71 | 485 | 556 |
|  | 7 | 0.01563 | 0.99224 | 4.69 | 33 | 188 | 221 |
|  | 8 | 0.00542 | 0.99766 | 1.63 | 13 | 64 | 77 |
|  | 9 | 0.00167 | 0.99933 | 0.50 | 5 | 19 | 24 |
|  | 10 | 0.00046 | 0.99979 | 0.14 | 1 | 5 | 6 |
|  | 11 | 0.00012 | 0.99991 | 0.04 | 0 | 1 | 1 |
| | 12 | 0.00003 | 0.99994<br>($\approx 1.0$) | 0.01 | 0 | 0 | 0 |
| Total | | 0.99994<br>($\approx 1.0$) | | 299.99<br>($\approx 300$) | 832 | 13267 | 14099<br>($\approx 14100$) |

The variability in X, in the Poisson probability distribution of variable X, is given by the variance σ^2^ = µ, where σ is the sample standard deviation. Here, standard deviation σ = µ^1/2^ = 2.773^1/2^ ≈ 1.665 (≈ 2), volunteers testing negative per sample.

From Table 1, the variable X for the cumulative probability of 0.93716 (or 93.7 % ≈ 95%) is: X = 5. It corresponds to X = µ + σ = 2.773 + 1.665 = 4.438 (≈ 5). Since for X = 0, there are no (0, zero) volunteers testing negative to vaccine, one may adopt the variable X to be significant within µ ± σ = 2.773 ± 1.665; that is, 1.108 to 4.438 (or 1 to 5). The variables X = 0, 1, 2, 3, 4 and 5 form group of volunteers testing negative to vaccine for 95% cumulative probability.

In mathematical statistics, the significance of variable X = 5 is the statement of assurance (certainty) that “not more than” 5 volunteers shall be testing negative to vaccine. The probability that a sample shall “fail to meet” the assurance is

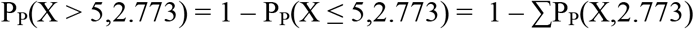

where the summation is from X = 0 to X = 5. From Table 1, ∑P_P_(X,2.773) = 0.93716 (≈ 0.95), implying P_P_(X > 5,2.773) = 0.06284 (≈ 5%), for 95% cumulative probability.

From Table 1, the number of samples with no volunteer testing negative, all volunteers testing positive, is 18.74, by chance. The corresponding percentage of samples is: (18.74 / 300) x 100 ≈ 6%.

Also, from Table 1, the number of samples with volunteers testing negative, corresponding to variable X = 1, 2, 3, 4 and 5 within ± σ, is 262.4 (≈ 262). It is about 88% of total 300 samples. Alternately, from Table 1, the corresponding number of volunteers testing negative is 709. It is about 85% of average 832 volunteers testing negative. Thus, on average, 87% of samples have average number of 2.773 (≈ 3, µ) volunteers testing negative, by chance. It implies that, on average, 724 (= 832 × 0.87) volunteers test negative in groups of 3 volunteers in 300 samples, by chance.

Also, from Table 1, the number of samples with volunteers testing negative, corresponding to variable X = 6, 7 and 8, is 18.16. It is about 6% of the total 300 samples. Alternately, from Table 1, the corresponding number of volunteers testing negative is 117. It is about 14% of the average 832 volunteers testing negative. Thus, on average, 10% of samples have 6 - 7 - 8 volunteers testing negative, by chance. It implies that, on average, 83 (≈ 832 × 0.10) volunteers test negative in groups of 7 in 300 samples, by chance.

A summary of groups of volunteers, average number of samples (%) and average number of volunteers, testing negative to vaccine in groups of 3 and 7 volunteers, is given in Table 2. On average, about 87% and 10% of samples have volunteers testing negative in groups of 3 volunteers and 7 volunteers, respectively, by chance. About 0.5% of samples, corresponding to variable X = 9 and 10, have volunteers testing negative to the vaccine as isolated cases, by chance.

**Table 2.** Summary of groups of volunteers, average number of samples (%) and average number of volunteers, in 300 samples of 47 volunteers each, testing negative for 94.1% vaccine efficacy, by chance (average number of volunteers testing negative = 832)

|  | Number of<br>volunteers testing<br>negative | Group of<br>volunteers<br>testing negative | Average number<br>of samples (%) | Average<br>number of<br>volunteers<br>testing negative |
| --- | --- | --- | --- | --- |
|  | 0 | 0 | 0%<br>(6%, all<br>volunteers<br>testing positive) | 0 |
| | 1 - 5 | 3 | 87% | $832 \times 0.87$<br>($\approx 724$) |
| | 6 - 7 - 8 | 7 | 10% | $832 \times 0.10$<br>($\approx 83$) |
| | 9 - 10<br>(Isolated cases) | - | 0.5% | $832 \times 0.005$<br>($\approx 4$) |
| Total | | | 103.5%<br>($\approx 100\%$) | 811<br>($\approx 832$) |

To summarize, on average, about 87% of samples, with volunteers testing negative within µ ± σ ≈ 3 ± 2, have groups of 3 volunteers testing negative. The standard deviation (σ) of the Poisson probability distribution is given by σ = µ^1/2^ = 1.665 ≈ 2 volunteers testing negative per sample. About 10% of samples have groups of 7 volunteers testing negative. By chance, about 0.5% of samples have volunteers testing negative to the vaccine as isolated cases.

The 95% confidence interval for the mean (µ = 2.773) of the sample for the Poisson probability distribution of variable X may be estimated from a χ^2^ table; for instance, from Reference [17, Table 26.8, page 984]. The 95% confidence limits for µ are given by

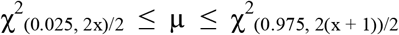

where χ^2^_0.025_ and χ^2^_0.975_ are the 2.5th percentile and 97.5th percentile, respectively, of the χ^2^ distribution. The discrete variable x refers to degrees of freedom (ν); x = 0 for a set of observations of a sample. The requisite limits of µ = 2.773 are obtained, with x = 0, as: 0 ≤ µ ≤ 3.689 (≈ 0 ≤ µ ≤ 4). (In Reference 17, the Table heading for “Q” for the lower limit of 2.5th percentile is: 1 – 0.025 = 0.975. The heading for “Q” for the upper limit of 97.5th percentile is: 1 – 0.975 = 0.025. ν = (x + 1). Thus, x = 0 is ν = 1, and x = 1 is ν = 2, etc. χ^2^ = 2µ, µ = χ^2^/2.) It implies that the range of the population mean µ, with 95% certainty, is: 0 ≤ µ ≤ 4.

The Moderna’s vaccine efficacy, for the confidence interval of 95% of the Poisson distribution, is in the range 89.3% - 96.8% of the population. The limits 89.3% and 96.8% mean that the probability (p) of negative response of vaccine is 0.107 and 0.032, respectively. Thus, the mean µ (µ = 47 x p) of the Poisson probability distribution of variable X per sample is 5.029 and 1.504, respectively. The mean µ = 1.504 and µ = 5.029 are within µ ± σ (= 2.773 ± 1.665) of the Poisson probability distribution of the variable X, with 95% certainty.

### 4.2 Statistical Analysis of Oxford’s 62.1% Vaccine Efficacy

In Phase 3, in Oxford’s vaccine testing procedure, there are 4807 volunteers at 19 centers in the UK and 4088 volunteers at 6 centers in Brazil; total of 8895 volunteers at 25 centers [8]. There are 8895 volunteers at effectively 75 (3 × 25 = 75) centers at an average of 118.6 (≈ 120) volunteers per center. (The number of groups of persons based on the age criterion is 3 at 25 centers.) The present study considers the vaccine testing in a population of 8895 volunteers, as N = 75 samples of n = 118.6 volunteers each. The vaccine efficacy is 62.1%. Thus, the probability (p) of negative response of vaccine in a sample is 37.9%; p = 0.379. Also, the average number of volunteers (µ) in a sample testing negative to vaccine is: µ = n x p = 118.6 × 0.379 = 44.949 = 45.0. The average number of volunteers testing negative is: µ × 75 = 45.0 × 75 = 3375 volunteers testing negative. (Alternately, the average number of volunteers testing negative is 3371, p × 8895 = 0.379 × 8895 = 3371.2 (≈ 3375), in the population of 8895.) Since p is finite, the Gaussian (or normal) probability distribution describes the vaccine testing procedure. (The Gaussian distribution function describes the data that is symmetric about the mean [6, Chapter 3]). The Gaussian distribution function, with mean equal to 45 (µ) volunteers testing negative to vaccine, is used to compute the probability of groups of volunteers (say, a group of 45 volunteers) testing negative to the vaccine in a sample. Hence, by chance, any groups of volunteers in 75 samples testing negative to vaccine are determined. These groups of volunteers testing negative for 95% confidence interval are examined vis-à-vis the average number of volunteers obtained with 41.0% and 75.7% vaccine efficacy (95% confidence interval 41.0% - 75.7%).

The Gaussian distribution function is given by

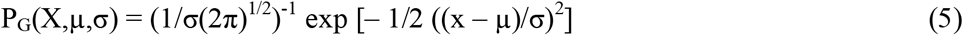

where X is the “continuous” random variable, µ the mean (average) number of volunteers per sample testing negative to vaccine and σ the sample standard deviation of variability in variable The variable X, in essence, depicts a range of number of volunteers testing negative to vaccine. The continuous function P_G_(X,µ,σ) is the probability density such that dP_G_(X,µ,σ) is the probability of X (number of volunteers) within an infinitesimal interval dX, around X: dP_G_(X,µ,σ) = P_G_(X,µ,σ) dX. The integral (area) of the distribution function P_G_(X,µ,σ), A_G_(X,µ,σ), between the limits µ ± zσ, is the probability that X (number of volunteers) deviates from µ less than zσ; z (= |(X – µ)| / σ)) and A_G_(z = ∞) = 1 [6, Chapter 3]. The area under the curve A_G_(X,µ,σ) versus z is tabulated elsewhere [Reference 6, Table C-2, page 308]. The table of A_G_(X,µ,σ) versus z is a universal table, for any set of values of (µ, σ). For instance, the probability of X (number of volunteers), around µ, between ± σ is the area 68.3% (≈ 68%). N x A_G_(X,µ,σ), N = 75, is the number of samples having a range of X (number of volunteers) testing negative to vaccine, by chance.

Since the sample size n = 118.6 is large (n > 100), the variability in variable X is given by the variance σ^2^ = np(1 – p), where σ is the sample standard deviation. (σ = (npq)^1/2^, q = 1 – p, for the binomial probability distribution.) Here, standard deviation σ = (118.6 × 0.379 × 0.621)^1/2^ = 5.28 = 5.3 (≈ 5), volunteers testing negative per sample.

(a) The mean µ is significant within ± σ. Thus, variable X (number of volunteers) lies within µ ± σ = 45.0 ± 5.3; that is, 39.7 (≈ 40.0) to 50.3 (≈ 50). Thus, on average, 68% (≈ 68.3%) of volunteers per sample have 45 - 50 volunteers (average number of 45 volunteers, µ) testing negative, by chance. On average, 51 (≈ 75 × 0.683 = 51.2) samples have 45 volunteers negative, by chance. The percentage of samples is: 51.2 / 75 = 68.3% (≈ 68%). The number of volunteers testing negative is 2295 (= 51 × 45); alternately, 3375 × 0.68 = 2295.

(b) Consider that variable X (number of volunteers) lies within ±1.5σ, around µ, µ ± 1.5σ = 45.0 ± 8; that is, 37 to 53. The area between ±1.5σ is 86.6% (≈ 87%). The difference of area between ±1.5σ and ±1σ is: 86.6% – 68.3% = 18.3% (≈ 18%). Thus, on average, 18% of volunteers per sample have (i) 37 - 40 volunteers (average number of 39 volunteers) and (ii) 50 - 53 volunteers (average number of 52 volunteers) testing negative, by chance. (The range of volunteers is obtained by combining with the range of volunteers in (a).) The probability of each group of volunteers in (i) and (ii) is 9%. On average, 7 (≈ 75 × 0.09 = 6.8) samples have (i) 39 volunteers testing negative and (ii) 52 volunteers testing negative, each, by chance. The percentage of samples in each case is 9% (6.8 / 75 = 9%). The number of volunteers testing negative is 637 (= 273 + 364); (i) 273 (= 7 × 39) and (ii) 364 (= 7 × 52).

(c) Consider that variable X (number of volunteers) lies within ±1.96σ (≈ ±2σ), around µ, µ ± 1.96σ = 45.0 ± 10.4; that is, 34.6 (≈ 35) to 55.4 (≈ 55). The area between ±1.96σ is 95%. The difference of area between ±1.96σ and ±1.5σ is: 95% – 86.6% = 8.4% (≈ 8%). Thus, on average, 8% of volunteers per sample have (i) 35 - 37 volunteers (average number of 36 volunteers) and (ii) 53 - 55 volunteers (average number of 54 volunteers) testing negative, by chance. (The range of volunteers is obtained by combining with the range of volunteers in (b).) The probability of each group of volunteers in (i) and (ii) is about 4%. On average, 3 (75 × 0.04 = 3) samples have (i) 36 volunteers testing negative and (ii) 54 volunteers testing negative, each, by chance. The percentage of samples in each case is: 3 / 75 = 4%. The number of volunteers testing negative is 270 (= 108 + 162); (i) 108 (= 3 × 36) and (ii) 162 (= 3 × 54).

A summary of average groups of volunteers, average number of samples (%) and average number of samples, of volunteers testing negative to vaccine in 75 samples is given in Table 3.

**Table 3.** Summary of average groups of volunteers, average number of samples (%) and average number of samples, in 75 samples of 120 volunteers each, testing negative for 62.1% vaccine efficacy, by chance

|  | Range of number of volunteers testing negative | Average group of volunteers testing negative | Average number of samples (%) | Average number of samples |
| --- | --- | --- | --- | --- |
|  | 35 - 37 | 36 | 4% | 3 |
|  | 37 - 40 | 39 | 9% | 7 |
|  | 40 - 50 | 45 | 68% | 51 |
|  | 50 - 53 | 52 | 9% | 7 |
|  | 53 - 55 | 54 | 4% | 3 |
| Total | | | 94%<br>($\approx 95\%$) | 71 |

The average number of samples for 95% probability is 71, which is less than the total number of samples of 75. The number of volunteers testing negative below 35, and above 55, is described by a random fluctuation; the probability is 0.025 (2.5%).

To summarize, on average, about 68% of samples, with volunteers testing negative within µ ± σ ≈ 45 ± σ ≈ 45 ± 5, have groups of 45 volunteers testing negative. The standard deviation (σ) of Gaussian probability distribution is given by σ ≈ (npq)^1/2^, q = 1 – p; σ = (118.6 × 0.379 × 0.621)^1/2^ = 5.28 = 5.3 (≈ 5), volunteers testing negative per sample. About 18% of samples, with volunteers testing negative within {(µ ± 1.5σ) – (µ ± σ)} have groups of 39 volunteers and 52 volunteers testing negative. The percentage of samples in each case is 9%. About 8% of samples, with volunteers testing negative within {(µ ± 1.96σ) – (µ ± 1.5σ)} have groups of 36 volunteers and 54 volunteers testing negative. (The area between µ ±1.96σ is 95%.) The percentage of samples in each case is 4%.

For the Gaussian probability distribution of variable X, the 95% confidence limits of the population mean (µ) based on a single sample, are calculated as:

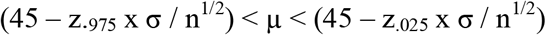

where the sample standard deviation σ = 5.3 and n^1/2^ = 118.6^1/2^ = 10.89 = 10.9 (≈ 11). The 2.5th percentile and 97.5th percentile limits of the Gaussian probability distribution, in terms of the variable z (= |(X – µ)| / σ)), are: z_.025_ = –1.96 and z_.975_ = 1.96. It implies that the range of population mean µ, with 95% probability, is: 44 < µ < 45.

The Oxford’s vaccine efficacy, for the confidence interval of 95%, is in the range 41.0% - 75.7% of the population. The limits 41.0% and 75.7% mean that the probability (p) of negative response of vaccine is 0.59 and 0.243, respectively. Thus, the average number of volunteers testing negative per sample (µ = 188.6 x p) is about 29 and 70, respectively. These average numbers of volunteers, testing negative to vaccine, are not within 35 to 55 (µ ± 1.96σ) of the Gaussian distribution of the variable X, with 95% certainty.

## 5. Conclusion

The three basic aspects of COVID-19 are presented in the present study.

The occurrence of coronavirus infection in human beings is statistically analyzed in terms of the number of coronavirus infected alveolar cells compared to normal alveolar cells in human lungs. The mole concept, that is, one mole of a substance (biological, virus, or vaccine) contains Avogadro’s number of particles (cells, viral particles, or specific antigens) is applied. It leads to the number of normal alveolar cells of size parameter 8000 nm per human lung volume as 10^9^. The number of coronavirus infections in terms of infected alveolar cells is estimated from the published Lower Respiratory Tract (LRT) load data in patients from Germany and China. The number of infected alveolar cells of size 100 nm per human lung volume is 10^3^. The number of coronavirus infections in infected alveolar cells (per lung volume) is much less than the number of normal alveolar cells (per lung volume). The Poisson probability distribution is aptly used to imply the appearance of symptoms, the incubation period, of coronavirus infection to be within day-3 to day-7of the exposure to infection. The cumulative probability, with mean incubation on day-5, is 75%. The cumulative probability, with mean incubation on day-5, for the infection to be between day-0 to day-10 of the exposure to infection is 98%.

Three vaccines to combat COVID-19, which adopt distinct paradigms while preparing them, are analyzed. These are Moderna’s mRNA-1273, Oxford-AstraZeneca’s ChAdOx1 nCoV-19 (AZD1222) and Bharat BioTech’s COVAXIN. The vaccine mRNA-1273 is a messenger (m) RNA-based vaccine that carries the SARS CoV-2 virus spike immunogen. The vaccine AZD1222 is a genetically modified preparation from chimpanzee adenovirus. The vaccine COVAXIN is prepared using live but inactivated SARS-CoV-2 virus. The mole concept leads to the antigen mass density per dose of each of these vaccines as 10 g cm^-3^, 0.1 g cm^-3^ and 1 g cm^-3^, respectively. Thus, the estimated mass density of an antigen dose is ∼ 10^0±1^ g cm^-3^. The vaccines are deemed to be compatible to neutralize the infection.

A statistical analysis is performed of the Moderna’s vaccine efficacy of 94.1% and Oxford’s vaccine efficacy of 62.1% in terms of groups of volunteers testing negative to the vaccine, by chance, for 95% certainty. The Moderna’s mRNA-1273 vaccination testing scenario comprises 300 samples of 47 volunteers each. The Moderna’s 94.1% vaccine efficacy, probability p = 0.059 (5.9%), implies that on average 3 volunteers per sample have a negative response to vaccine, by chance. Since the probability of negative response of vaccine is small, the Poisson probability distribution for 95% cumulative probability is used to describe the vaccination testing. Thus, 87% of samples (i.e., 261 samples) have groups of 3 volunteers testing negative to the vaccine. Also, 10% of samples (i.e., 30 samples) have groups of 7 volunteers testing negative to vaccine. About 0.5% of samples (i.e., 2 samples) have groups of 9 - 10 volunteers testing negative to vaccine, with a small probability. About 6% of samples (i.e., 18 samples) have all volunteers testing positive to vaccine. (The total number of samples is 311, which is roughly 300 samples.) The Oxford’s ChAdOx1 nCoV-19 vaccination testing scenario comprises 75 samples of 120 volunteers each. The Oxford’s 62.1% vaccine efficacy, probability p = 0.379 (37.9%), implies that on average 45 volunteers per sample have a negative response to vaccine, by chance. Since the probability of negative response of vaccine is finite, the Gaussian (or normal) probability distribution for 95% probability is used to describe the vaccine testing. Thus, 68% of samples (i.e., 51 samples) have groups of 45 volunteers testing negative to vaccine. Also, 9% of samples (i.e., 7 samples) of groups of 39 volunteers, and another 9% of samples (i.e., 7 samples) of groups of 52 volunteers, test negative to the vaccine. Additionally, 4% of samples (i.e., 3 samples) of groups of 36 volunteers, and another 4% samples (i.e., 3 samples) of groups of 54 volunteers, test negative to vaccine. No sample has all volunteers testing positive to vaccine. (The total number of samples is 71. In addition, about four samples may have a random fluctuation of groups of volunteers below 35 volunteers or above 55 volunteers, with a low probability.) A generalized vaccination testing procedure is one in which both sample size of volunteers being vaccinated and number of samples is 100, each, in order to evaluate vaccine efficacy. It makes the total number of samples 200, including the placebo group of volunteers.

The statistical study of the occurrence of coronavirus infection implies that the 10-day quarantine be adopted to isolate an individual as a precautionary measure. The compatibility study of the vaccines implies that the mixed doses of the vaccines may be used to neutralize the infection. The statistical analysis of the testing procedure provides, in general, the probabilistic distribution of samples of volunteers testing negative to the vaccine. A vaccine, irrespective of its efficacy being high or low, is necessary for mass immunization to slow and prevent the spread of infection the COVID-19 pandemic.

## Acknowledgements

The authors gratefully acknowledge Dr. B. Madhava Reddy and Dr. P.N. Vijaykumar, formerly of the National Physical Laboratory, for their helpful comments.

